# The Effect of Famotidine on SARS-CoV-2 Proteases and Virus Replication

**DOI:** 10.1101/2020.07.15.203059

**Authors:** Madeline Loffredo, Hector Lucero, Da-Yuan Chen, Aoife O’Connell, Simon Bergqvist, Ahmad Munawar, Asanga Bandara, Steff De Graef, Stephen D. Weeks, Florian Douam, Mohsan Saeed, Ali H. Munawar

## Abstract

The lack of coronavirus-specific antiviral drugs has instigated multiple drug repurposing studies to redirect previously approved medicines for the treatment of SARS-CoV-2, the coronavirus behind the ongoing COVID-19 pandemic. A recent, large-scale, retrospective clinical study showed that famotidine, when administered at a high dose to hospitalized COVID-19 patients, reduced the rates of intubation and mortality. A separate, patient-reported study associated famotidine use with improvements in mild to moderate symptoms such as cough and shortness of breath. While a prospective, multi-center clinical study is ongoing, two parallel *in silico* studies have proposed one of the two SARS-CoV-2 proteases, 3CL^pro^ or PL^pro^, as potential molecular targets of famotidine activity; however, this remains to be experimentally validated. In this report, we systematically analyzed the effect of famotidine on viral proteases and virus replication. Leveraging a series of biophysical and enzymatic assays, we show that famotidine neither binds with nor inhibits the functions of 3CL^pro^ and PL^pro^. Similarly, no direct antiviral activity of famotidine was observed at concentrations of up to 200 μM, when tested against SARS-CoV-2 in two different cell lines, including a human cell line originating from lungs, a primary target of COVID-19. These results rule out famotidine as a direct-acting inhibitor of SARS-CoV-2 replication and warrant further investigation of its molecular mechanism of action in the context of COVID-19.

## INTRODUCTION

A large part of the current therapeutic discovery effort against the severe acute respiratory syndrome coronavirus 2 (SARS-CoV)-2 is focused on drug repurposing^1^. Of such agents, only remdesivir has thus far shown clinical evidence of antiviral effect^2^, while several others have not met their primary endpoints in various clinical studies^3,4^. Recently, famotidine has gained attention as a therapeutic option against SARS-CoV-2, initially based on anecdotal evidence of its positive effects in COVID-19 patients in China. Famotidine (PEPCID^®^), a histamine-2 receptor (H2R) antagonist, is an FDA approved drug for the treatment of gastroesophageal reflux disease (GERD) and gastric ulcers^5^.

Earlier reports of the beneficial effect of famotidine in China were recently supported by a retrospective clinical study involving 1620 patients in the U.S., which noted that hospitalized COVID-19 patients receiving a total median dose of 136 mg famotidine, in oral or IV formulation once daily, for 6 days had a reduced risk of death or intubation^6^. Another study involving 10 non-hospitalized patients linked the use of high-dose oral famotidine (240 mg per day for a median of 11 days) with patient-reported improvements in symptoms such as shortness of breath and cough^7^. These two reports conclude that the use of high-dose famotidine may be associated with improvements in both mild and severe symptoms of COVID-19. While a large, multi-center clinical trial to confirm these observations is in progress, the mechanism by which famotidine purportedly improves the clinical outcomes in COVID-19 patients is unknown. *In silico* modeling and molecular docking studies have separately suggested either of the two SARS-CoV-2 proteases as potential targets of famotidine activity^8,9^. In one computational study, Wu et. al. docked a library of approved drugs on to the available X-ray crystal structure of the 3-chymotrypsin-like protease (3CL^pro^) of SARS-CoV-2, identifying famotidine as one of the drugs likely to act on the protease ^8^. Other computational reports have predicted famotidine as an inhibitor of the Papain-like protease (PL^pro^), a second SARS-CoV-2 protease^9^. Together, these studies have raised the prospect of a direct antiviral effect of famotidine on SARS-CoV-2 replication. While both proteins are attractive targets for SARS-CoV-2 drug development^10–19^, there are at present no clinical-stage or approved drugs targeting either protein. The possibility of famotidine, an approved drug, acting on SARS-CoV-2 proteases is of significant clinical interest. In this study, we performed an array of biochemical, biophysical, and antiviral experiments to test if famotidine is an effector of SARS-CoV-2 proteases and whether it inhibits virus replication in cultured cells.

## MATERIALS AND METHODS

### Compounds

Famotidine was acquired from Sigma Aldrich (Missouri, USA; cat. No. F6889). **Compound 6,** a previously reported inhibitor of SARS-CoV-2 PL^pro^ function^20^ was acquired from MedChem Express, Inc. (New Jersey, USA; cat no. HY-17542). **ML188**, a compound with known 3CL^pro^ inhibitory activity^15^ was also acquired from MedChem Express, Inc. (cat. no HY-136259). Similarly, remdesivir. (cat. No HY-104077) an inhibitor of SARS-CoV-2 replication^2^ was purchased from the same vendor. All compounds were dissolved in 100% DMSO at 100mM.

### Cloning, expression, and protein purification

The complete sequences encoding 3CL^pro^ and residues 746-1060 of PL^pro^ (Wuhan-Hu-1 isolate, GenBank accession NC_045512) were cloned into a charge modified SUMO fusion expression vector, generated in-house. The fusion protein was expressed for 24 hours in Rosetta-2 (DE3) pLysS at 18°C in ZYP-5052 autoinducing media. Harvested cells were resuspended in 50 mM Hepes pH 7.5, containing 150 mM NaCl and lysed by sonication. The clarified supernatant was loaded onto a HiTrap HP SP column (Cytiva, Massachusetts, USA; cat no. 17115201) and the target fusion protein was captured in a cation-exchange chromatography step and eluted using a NaCl gradient. SUMO hydrolase was added to the pooled fractions to liberate the target protein and the sample dialyzed against 20 mM Tris, 10 % v/v glycerol, 5 mM DTT pH 7.0 overnight at 4°C. The protein was reloaded on the HiTrap HP SP column to remove the SUMO protein and hydrolase in a subtractive step. The flow-through, containing 3CL^pro^ or PL^pro^ was further purified by anion exchange chromatography using a HiTrap HP Q column (Cytiva; cat. no. 17115401) employing a NaCl gradient to elute the protein. Pooled fractions were further purified by size exclusion chromatography in 20 mM Tris pH 7.4, 150 mM NaCl and 5 mM DTT. The final protein was concentrated to 4 mg/mL for PL^pro^ and 5 mg/mL for 3CL^pro^ and flash frozen in aliquots.

### *In-vitro* viral enzyme assays

#### PL^pro^ proteolytic activity assay using ubiquitin-AMC

PL^pro^ activity was measured in a 384 well plate format (Corning #3574) in a kinetic assay using the fluorogenic substrate Ubiquitin-AMC (Boston Biochem, Inc. Massachusetts, USA; cat. No. U-550) with excitation and emission wavelengths of Ex355nm/Em:460nm. The protocol followed previously reported conditions with minor modifications^13,16^. Fluorescence was monitored at 25°C, every 5 min for 50 min in a Victor X5 (Perkin Elmer) multimode plate reader. Optimal enzyme and substrate concentrations were found to be 550 nM PL^pro^ titrating the substrate in the range of 0.2 – 3 μM. The assay buffer (20 μL) contained 25 mM HEPES pH 7.5, 100 mM NaCl, 0.1 mg/ml BSA, and 550 nM PL^pro^. The test inhibitor, famotidine and the PL^pro^ control inhibitor (**compound 6**) were both titrated in the concentration range of 0.01 μM – 200 μM. Compounds were incubated with the enzyme in the plate for 30 minutes at 25°C before the reaction was started by the addition of 1 μM Ub-AMC. All samples were run in triplicates and their initial slopes were converted from relative fluorescence units (RFU)/ min to μmol AMC/min using an AMC standard curve and plotted against compound concentrations tested.

#### 3CL^pro^ proteolytic activity assay

3CL^pro^ activity was assayed in a 384 well plate using the 3CL^pro^ FRET substrate (AnaSpec, Inc. California, USA; cat. no. AS-65599) with excitation and emission wavelengths of Ex: 490nm/Em: 535 nm. A previously reported protocol was used with some modifications^12,17^. The kinetics of fluorescence change were monitored every minute for 25 min. Optimal concentrations for 3CL^pro^ and substrate were 150 nM and 600 nM respectively. A previously reported 3CL^pro^ inhibitor^14^, **ML188**, was used as a positive control for inhibition, both control and test compounds were titrated in the concentration range of 0.01 μM – 200 μM. Initial slopes of RFU/min were converted to μM hydrolyzed substrate/ min using a standard curve of HiLyte Fluor488 amine, TFA salt.

#### Biochemical Data Analysis

After subtraction of background fluorescence readings, values of Km and EC50 were obtained by fitting the experimental data with the Michaelis-Menten (y=(Vmax*x)/(Km+x)) and the four parameters logistic (4PL) equations (y= min + (max-min)/(1+(x/EC50)^Hillslope)) respectively, using GraphPad Prism 8.

### Dynamic Scanning Fluorimetry

Thermal unfolding of proteins was monitored in a 20 uL volume in Micro-Amp EnduraPlate Optical 384-well Clear Reaction Plates (ThermoFisher: cat no. 4483285). Reactions contained 50 mM HEPES pH 7.5, 62.5 mM NaCl, 7 μM 3CL^pro^ or PL^pro^, 5% DMSO, and 4x SYPRO-orange protein gel stain (ThermoFisher Scientific, Massachusetts, USA; cat no. S6651). Famotidine and the positive controls **ML188** and **compound 6** for 3CL^pro^ and PL^pro^, respectively, were incubated with the protein for 15 minutes before the addition of SYPRO orange. Plates were covered with Micro-Amp Optical Adhesive Film (ThermoFisher Scientific, Massachusetts, USA; cat no. 4360954) and run on Applied Biosystems 7900HT (California, USA) real time PCR instrument. Samples were incubated at 25 °C for 2 min followed by an increase in temperature of 1°C/min up to 95°C. Fluorescence was monitored continuously. Each sample was run in triplicate and compounds were tested at 1 mM, 2.5 mM, and 5 mM. The melting temperature (*T_m_*) was obtained from the first derivative of the raw thermal denaturing data were determined and smoothed to calculate melting temperature (*T_m_*) values ^21^.

### Surface Plasmon Resonance (SPR)

SPR studies were performed on a Biacore 3000 instrument (Cytiva, Massachusetts, USA) at 10 °C. The PL^pro^ and 3CL^pro^ proteins were biotinylated by minimal biotinylation approach with the EZ-LINK Sulfo-NHS-LC-LC-biotin reagent (ThermoFisher Scientific, Massachusetts, USA; cat no. A3 53 58) and immobilized on a neutravidin coated CM5 sensor chip to a level of 4000 response units (RU). The protein used during immobilization was at 1 μM for PL^pro^ and 1μM for 3CL^pro^. During the course of the assay different concentrations of compounds were injected. The Compounds were serially diluted (2-fold) in a running buffer of 25 mM HEPES pH 7.0, 200 mM NaCl, 2 mM TCEP, 0.005% P20 and 1% DMSO. Famotidine, and the control inhibitors, **compound 6** and the **ML188** were tested up to a maximal dose of 100 μM, 50 μM and 5μM, respectively. The final response was obtained by subtracting the blank channel (without protein) and a buffer injection across the sample channel. Raw data were analyzed in the Scrubber2 program (BioLogic Software) by fitting the data to a simple 1:1 equilibrium and kinetic model.

### Antiviral assays

#### Viruses and titration

Virus infectivity assays were carried out using the 2019-nCoV/USA-WA1/2020 isolate of SARS-CoV-2 (NCBI accession number: MN985325), obtained from the Centers for Disease Control and Prevention and BEI Resources (Virginia, USA). The virus stock was propagated in Vero E6 cells and virus titers determined using plaque formation assays, as described previously^22^.

#### Antiviral assays

Human lung A549 cells expressing SARS-CoV-2 entry factors and African Green Monkey kidney Vero E6 cells were maintained in DMEM supplemented with 10% fetal bovine serum (FBS). The cells were seeded into poly-L-lysine-coated 96-well plates at a density of 15,000 cells per well. The cells were then treated for 4h with five-fold serial dilutions of famotidine, ranging between 0.32 μM and 200 μM. DMSO served as a negative control, while 5-fold serial dilutions of remdesivir, ranging between 0.1 μM and 62.5 μM, served as a positive control. The cells were then infected with SARS-CoV-2 at a multiplicity of infection (MOI) of 0.1. To infect cells, the compound-containing medium was removed, and the cells were incubated with the virus for 1h at 37°C. The virus inoculum was then removed, and the cell monolayer was rinsed twice with 1X PBS. The compounds were added back followed by incubation for 72h, after which the cell culture medium was harvested for quantitative real-time PCR (qRT-PCR) and plaque assays, while the cells were fixed with 4% paraformaldehyde for immunofluorescence microscopy.

#### Virus RNA extraction and quantitative real-time PCR (qRT-PCR)

RNA was isolated from the cell culture supernatant of SARS-CoV-2-infected cells using the Quick-RNA Viral Kit (Zymo, California, USA cat no. R1035) according to the manufacturer’s instructions. Viral RNA was quantified using single-step RT-quantitative real-time PCR using the qScript One-Step RT-qPCR Kit (Quantabio, Massachusetts, USA; cat no. 95058) with primers and Taqman probes targeting the SARS-CoV-2 E gene as previously described^23^. Data were acquired using a Quantstudio3 Real-Time PCR System (Applied Biosystems) using the following conditions: 55°C for 10 min, denaturation at 94°C for 3 min, 45 cycles of denaturation at 94°C for 15 sec, and annealing at 58°C for 30 sec. The primers and probe used were as follow: E_Sarbeco_Forward: ACAGGTACGTTAATAGTTAATAGCGT; E_Sarbeco_Probe: FAM-ACACTAGCCATCCTTACTGCGCTTCG-BBQ; E_Sarbeco_Reverse: ATATTGCAGCAGTACGCACACA. For absolute quantification of viral RNA, a 389 bp fragment from the SARS-CoV-2 E gene was cloned onto pIDTBlue plasmid under an SP6 promoter using NEB PCR cloning kit (New England Biosciences, Massachusetts, USA; cat no. E1202S). The cloned fragment was then *in vitro* transcribed using the mMessage mMachine SP6 transcription kit (ThermoFisher, Massachusetts, USA; cat no. AM1340) to generate the qRT-PCR standard.

#### Immunofluorescence microscopy

Virus-infected cells were fixed with 4% paraformaldehyde for 30 minutes. The fixative was removed, and the cell monolayer washed twice with 1X PBS. The cells were then permeabilized and stained with an anti-SARS-CoV Nucleocapsid (N) antibody (Rockland Inc., Pennsylvania, USA; cat. no. 200-401-A50; 1:2,000 dilution). Incubation with the primary antibody was performed overnight at 4°C. The cells were then washed 5 times with 1X PBS and stained with Alexa 568-conjugated anti-rabbit antibody (1:1000 dilution) in the dark at room temperature for 1h and counterstained with DAPI. Images were captured using EVOS M5000 Imaging System (ThermoFisher Scientific, Massachusetts, USA). Quantitation and analysis of the fixed cell images was carried out using the MuviCyte Live-Cell Imaging System (PerkinElmer, Massachusetts, USA). At least 7-10 microscopic fields were imaged per well using a 10X objective lens, the number of cells positive for the SARS-CoV-2 N protein and the nuclear DAPI stain, were counted. For each image, the percentage of DAPI-positive cells expressing the viral N protein were calculated, and the mean±SD of multiple images for each condition was plotted.

#### Cytotoxicity/Cell viability assay

The CellTiter-Glo Luminescent Cell Viability Assay (Promega, Wisconsin, USA; cat no. G7570) was used to determine the cytotoxic effects of the compounds. Briefly, the cells were incubated with five-fold serial dilutions of famotidine or remdesivir for 72h, after which the CellTiter-Glo Reagent was added to each well in a volume equal to the volume of the culture medium. The contents were mixed by shaking the plate on an orbital shaker for 2 min, followed by a 10 min incubation at room temperature. Luminescence was recorded using a Varioskan LUX multimode plate reader (ThermoFisher Scientific, Massachusetts, USA).

## RESULTS

### Famotidine is not an inhibitor of SARS-CoV-2 proteases

Processing of the SARS-CoV-2 polyprotein is critical to the generation of a functional virus replication complex^11,18,24^. To carry out this essential proteolytic function, the SARS-CoV-2 genome encodes two cysteine proteases, called PL^pro^ and 3CL^pro18^. Due to their critical roles in viral polyprotein processing and virus proliferation, both proteases are considered attractive targets for drug discovery^10,11,13–17,20^. Since *in silico* docking studies have predicted these proteases as putative molecular targets of famotidine^6,8,9^, we methodically investigated the effect of famotidine on the catalytic functions of each protease.

First, we developed an *in vitro* activity assay of PL^pro^. PL^pro^ is a protease domain found within the large multi-domain nsp3 protein encoded by SARS-CoV-2. While many coronaviruses encode two papain-like proteases, SARS-CoV, MERS-CoV and SARS-CoV-2 possess only one PL^pro^, which processes the amino-terminal end of the viral polyprotein liberating nsp1, nsp2 and nsp3 ^19,20^. Additionally, PL^pro^ deubiquitinates host cell proteins by cleaving the consensus motif of LXGG^18,19^ and is known to efficiently hydrolyze both diubiquitin and synthetic peptide substrates^19^. We leveraged the deubiquitinating property of PL^pro^ to set up a functional activity assay using ubiquitin-AMC, a fluorogenic substrate cleavable by PL^pro^. Upon incubation with PL^pro^, the ubiquitin is recognized and cleaved at the C-terminus to liberate the AMC (amido-4-methylcoumarin) fluorophore which results in increased fluorescence that is read using excitation and emission wavelengths of 355/460 nm. We assessed the ability of famotidine to inhibit the proteolytic activity of PL^pro^ at a broad range of drug concentrations vis-à-vis **compound 6**, a previously reported inhibitor of PL^pro^ activity^20^. Experimental conditions including protein and substrate concentrations, buffer composition, and assay kinetics were optimized using **compound 6**. While **compound 6** inhibited PL^pro^ activity with the expected low single-digit μM IC_50_ values, famotidine showed no reduction in PL^pro^ activity in the titrated range of 0.01-200 μM (Figure 1A).

**Figure 1.**
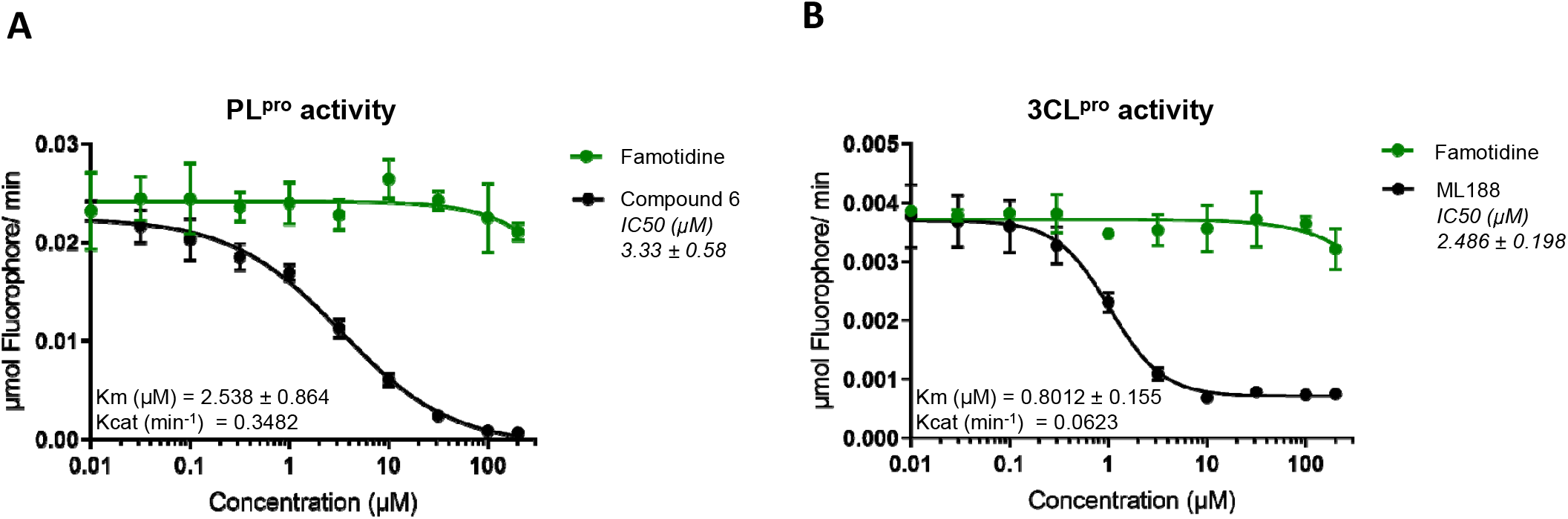
Effects of famotidine on PL^pro^ and 3CL^pro^ protease activity. *In vitro* inhibition assays (IC_50_) of PL^pro^ **(A)** and 3CL^pro^ **(B)** activity show that famotidine had no effect on either of the two SARS-CoV-2 proteases. IC_50_ values represent inhibition of viral protease activity by control compounds (black) and/or famotidine (green) when tested at various concentrations. The initial slopes of protein catalytic activity were converted from RFU/min to nmole fluorophore/min. Values are mean ± standard deviation of triplicates. The compounds tested in this experiment neither quenched fluorescence nor produced auto-fluorescence.

We next tested whether famotidine can inhibit the enzymatic activity of 3CL^pro^, the second protease encoded by the SARS-CoV-2 genome. This protein, also referred to as the main protease (M^pro^) or nsp5, cleaves the viral polyprotein at 11 unique sites^11^. This proteolytic activity generates multiple individual functional proteins required for the assembly of the SARS-CoV-2 replication/transcription complex, which drives viral genome replication^24^. Owing to its central role in the coronavirus life cycle, 3CL^pro^ has received significant attention as a drug target resulting in the discovery of several potent inhibitors ^10,14,15,17^ Native 3CL^pro^ exists as a homodimer and requires dimerization for its proteolytic activity^11^. The catalytic mechanism of 3CL^pro^ activity is typical of cysteine proteases, where the Cys-His catalytic dyad drives sitespecific cleavage of substrates. We evaluated the enzymatic activity of 3CL^pro^ using a FRET-peptide substrate that quenched fluorescence in its intact form, however, cleavage of the peptide substrate by 3CL^pro^ produced fluorescence that could be measured at the excitation/emission wavelengths of 490/535 nm. The inclusion of **ML188**, a previously reported 3CL^pro^ inhibitor served as a control, also aiding assay setup and optimization. Results of the FRET assay for various **ML188** and famotidine concentrations are shown in Figure 1B. Both drug compounds were tested between a range of 0.01-200 μM. While **ML188** produced a dose-dependent inhibition of 3CL^pro^ activity with an expected IC_50_ of 2.4 μM, famotidine did not inhibit 3CL^pro^ activity. These two experiments indicate that famotidine does not interfere with the catalytic activity of either of the two SARS-CoV-2 proteases.

### Famotidine does not directly engage PL^pro^ or 3CL^pro^ of SARS-CoV-2

The function of many enzymes, such as proteases and kinases, can extend beyond their catalytic roles and includes a wide spectrum of non-catalytic activities such as allosteric regulation, scaffolding, protein-protein interactions, and protein-DNA interactions^25^. To rule out whether famotidine could bind away from the active site of the two viral proteases, and exert an effect through interference with non-proteolytic functions, we asked if famotidine is able to bind directly with either of the two SARS-CoV-2 proteases. For this, we employed two distinct biophysical techniques i.e. surface plasmon resonance (SPR) and differential scanning fluorimetry (DSF), that are routinely used to probe drug-protein engagement.

For our SPR studies, the biotinylated viral proteases were captured to a high density on sensor chips via neutravidin, permitting real-time detection of small-molecule binding to the target viral proteases. Engagement of the small-molecule compounds was recorded as an increase in dose-dependent response units (RU) during the assay. Experimental conditions including buffer composition and temperature were optimized using the control compounds prior to conducting the famotidine studies. The equilibrium dissociation constant (*K_d_*) values were determined using both a kinetic analysis and fit to a binding isotherm of the dose response data (Figure 2). The observed *K_d_* values for the known 3CL^pro^ and PL^pro^ inhibitors (Supplementary Information <u>Table S1</u>) were consistent with the published IC_50_ data^14,20^ indicating the robustness of our assay methodology. Under these optimized conditions, famotidine was not found to interact with either of the two viral proteases at concentration ranges of up to 100 uM.

**Figure 2.**
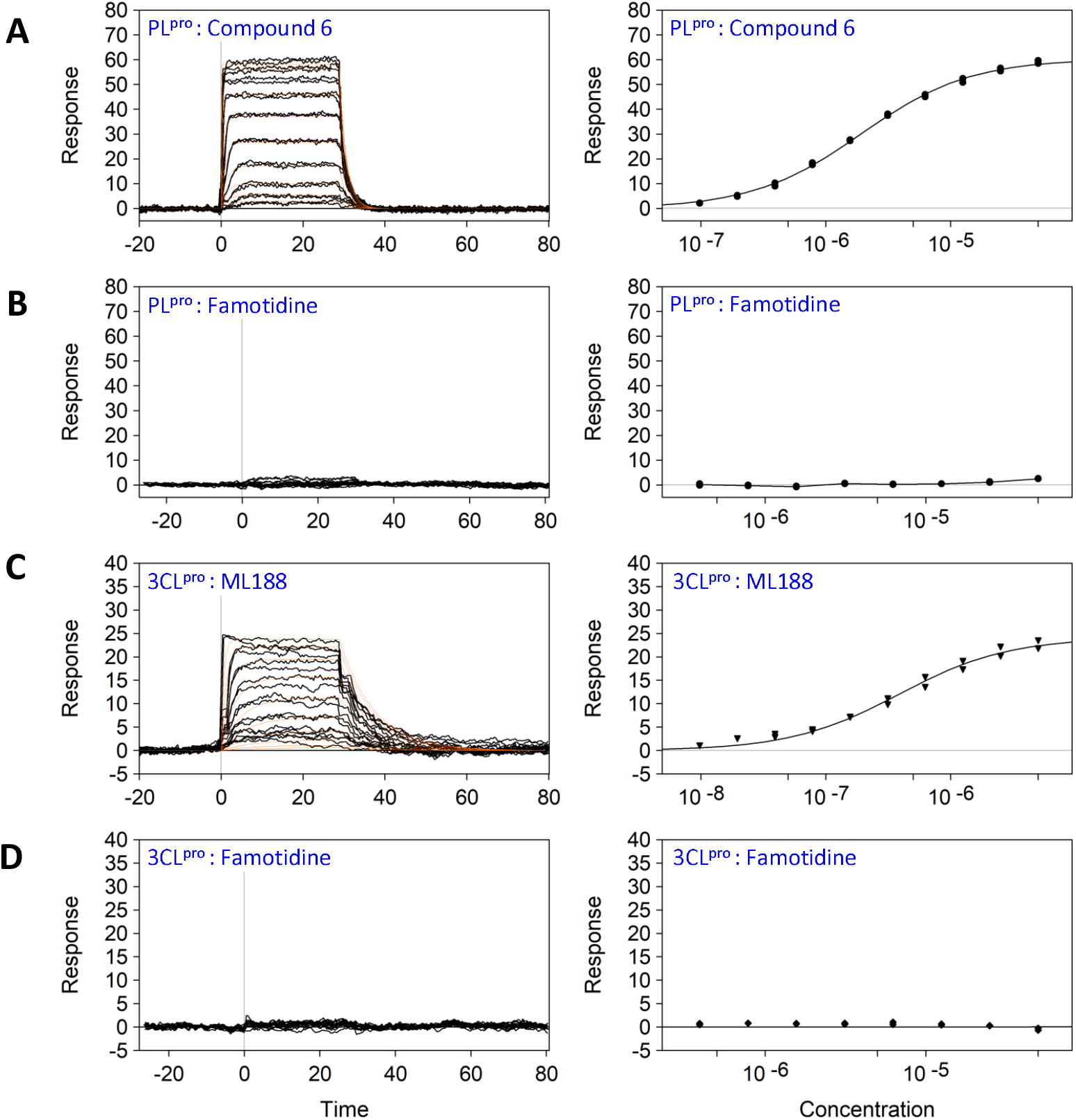
Binding of famotidine to PL^pro^ and 3CL^pro^ analyzed by SPR. Soluble biotinylated PL^pro^ (**A & B**) and 3CL^pro^ (**C & D**) were immobilized on a neutravidin-coated sensor chip and a range of compound concentrations were injected with solvent (DMSO) corrections. Both, **(A)** *compound 6,* the known PL^pro^ inhibitor and **(C)** *ML188,* the 3CL^pro^ inhibitor displayed dose-dependent binding to PL^pro^ and 3CL^pro^, respectively. Whereas, **(B and D)** no binding of famotidine was detected to either protein. The dissociation constant *(K_d_)* values for the control compounds are shown in Table 1 (supplementary materials).

To validate the results obtained from our SPR analysis, we employed an orthogonal DSF assay. DSF is a fluorescence-monitored thermal denaturing technique in which the melting temperature (*T_m_*) of a protein is tracked via fluorescence as the sample temperature is incrementally raised in the presence of a hydrophobic dye. Drug binding to its target protein is known to stabilize (or destabilize) protein structure resulting in a variation of *T_m_* profiles in the absence or presence of a drug. DSF provides definitive confirmation of target engagement as the increase in thermal unfolding temperature (Δ*T_m_*) is only achieved when the compounds bind to the folded state of the protein. The *ΔT_m_* is proportional to the *K_d_* of the interaction and concentration of the compound. We tested the ability of famotidine and the control inhibitors to alter the thermal stability profiles of PL^pro^ and 3CL^pro^. An optimal signal profile was obtained with 7μM PL^pro^ or 3CL^pro^. Both proteins were tested separately in the presence of DMSO (-ve control), their respective control inhibitors, and famotidine at concentrations of 1, 2.5 and 5mM. In agreement with the SPR data, the control inhibitors produced a quantitative increase in observed *T_m_* (Figure 3). While **compound 6**, the known PL^pro^ inhibitor, stabilized PL^pro^ by a *T_m_* of 5.5ūC (Figure 3A), and **ML188**, the 3CL^pro^ inhibitor, produced a *T_m_* shift of 4.8 C (Figure 3B), famotidine did not alter the *T_m_* of either of the two viral proteases. Taken together, the biophysical data decisively rules out the possibility of famotidine exerting its effect on PL^pro^ or 3CL^pro^ through interference with catalytic or non-catalytic protein functions as it is unable to bind with either of the two proteases.

**Figure 3.**
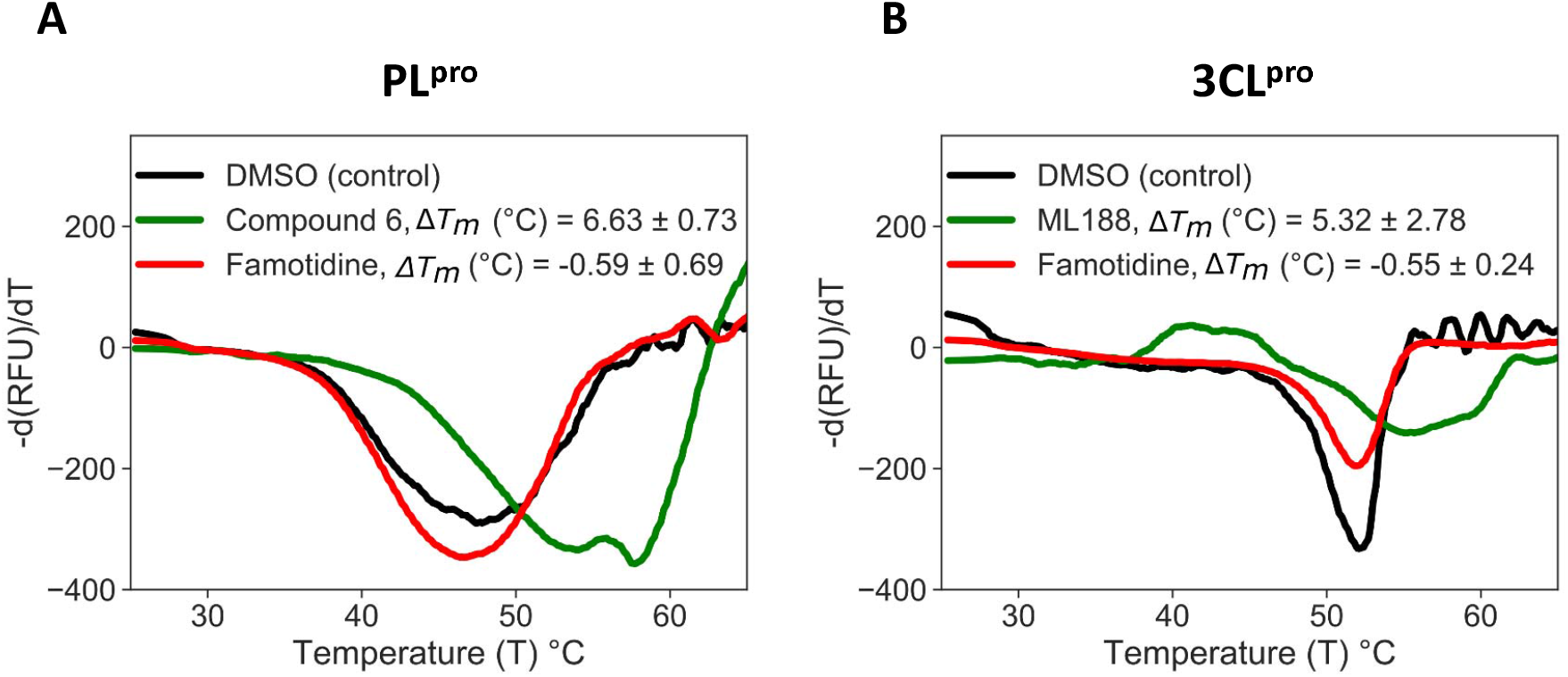
DSF assays of famotidine binding to PL^pro^ and 3CL^pro^. Fluorescence-monitored thermal denaturation assay showing the melting curve (first-derivative of dissociation) for each of the two proteins (7μM) in the presence or absence of compounds (2.5mM). **(A)** PL^pro^ melting curves for DMSO control (black), *compound 6* (green) and famotidine (red) show that while *compound 6* stabilizes the protein *ΔT_m_* by over 6.5°C, famotidine is unable to shift the *ΔT_m_*. Similarly, in **(B)** while *ML188* (green) stabilizes 3CL^pro^ *ΔT_m_*, by 5.3 °C, famotidine (red) does not shift the melting temperature of 3CL^pro^. The values are mean ± standard deviation of three independent replicates.

### Famotidine does not inhibit SARS-CoV-2 replication in cultured cells

Having established that famotidine does not inhibit SARS-CoV-2 proteases, we investigated the ability of famotidine to block virus replication in cell culture. For this, we infected Vero E6 cells, a commonly used cell model of SARS-CoV-2 infection derived from the African green monkey kidney. Infection efficiency was quantified through multiple, orthogonal readouts, including quantitative real-time PCR (qRT-PCR), plaque formation, and immunofluorescence. Remdesivir inhibited viral replication with an estimated half-maximum inhibitory concentration (IC_50_) value of 3.3 μM, as determined by immunofluorescence (Figure 4A). In contrast, famotidine did not produce any measurable inhibition at concentrations of up to 200 μM at 72h post infection. Similar results were obtained when viral replication was examined by infectious virion production using plaque formation assays or by quantifying viral RNA copy numbers in the cell culture medium using qRT-PCR (Supplementary Figures S1 and S2).

**Figure 4.**
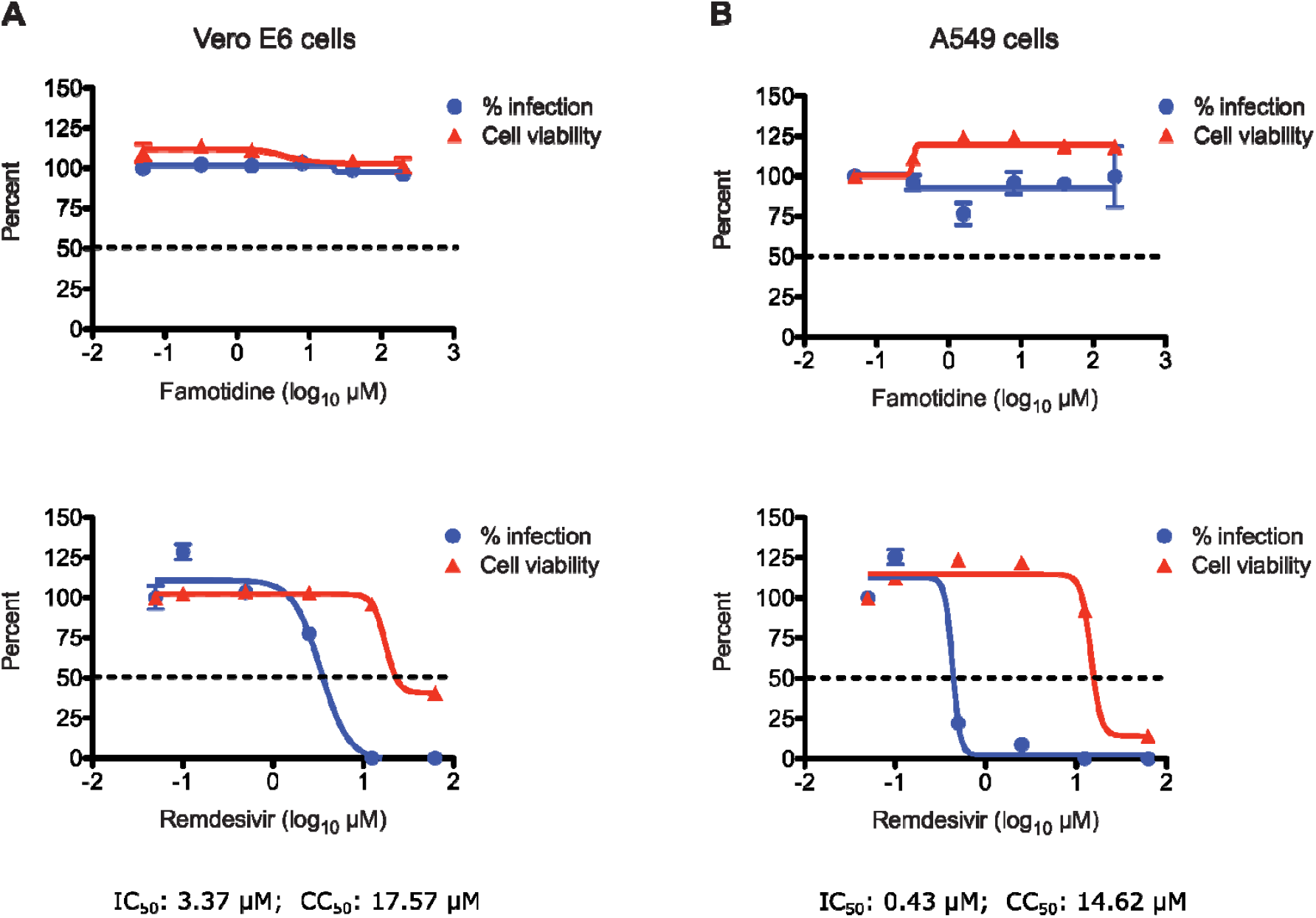
Antiviral activity of famotidine in Vero E6 and human lung A549 cells. Percent inhibition of SARS-CoV-2 replication and cytotoxicity are shown in the presence of a range of famotidine (top) and remdesivir (bottom) concentrations for **(A)** Vero E6 cells and **(B)** human lung A549 cells. IC_50_ values (blue) represent the antiviral activity of the drug compounds and CC_50_ values (red) represent cytotoxicity of the drugs. Infection was assessed through quantitation of virus-treated cells that stained positive for the viral nucleocapsid protein, 72 hours post infection. Cell viability of the corresponding compound concentrations on the cells was measured using the CellTiter-Glo assay. Values reported are mean ± standard deviation of triplicates.

To confirm these results in a more physiologically relevant cell model of SARS-CoV-2 infection, we assessed the antiviral activity of famotidine in human lung A549 cells. These cells were engineered to express essential SARS-CoV-2 entry factors, ACE2 and TMPRSS2. The cells were infected with SARS-CoV-2 and cultured in the absence or presence of the control or test compounds. Virus replication (infection) efficiency was measured and reported as a function of compound concentration (Figure 4). While remdesivir strongly inhibited virus replication in a dose-dependent manner with an IC_50_ value of 0.43 μM, famotidine had no measurable effect (Figure 4B). Our results are consistent with previously reported studies in which remdesivir exerted a greater antiviral effect in human lung A549 cells than in Vero E6 cells^26^.

In-parallel cytotoxicity assays, carried out in both Vero E6 and A549 cells, showed that famotidine was not toxic up to the highest tested concentrations of 200 μM (Figure 4 A and B). Remdesivir, on the other hand, exhibited dose-dependent cytotoxicity at higher concentrations, well above its IC_50_. Together, these results show that famotidine does not inhibit SARS-CoV-2 replication in cultured cells and that its purported clinical benefit may be due to an alternative mechanism of action.

## DISCUSSION

Two *in silico* studies have separately predicted the 3CL^pro^ or PL^pro^ of SARS-CoV-2 as potential molecular targets of famotidine^8,9^, implying that famotidine associated improvement in COVID-19 patients may be due to a direct antiviral mechanism of action^6^. Despite recent advances in computational techniques, there are several challenges associated with the use of molecular docking to predict protein-ligand interactions accurately. Some of these challenges arise from the flexibility of the target protein, lack of prior knowledge of drug-binding sites, and protonation states of target amino acids. While results obtained from molecular docking can serve as a basis for new hypotheses, experimental validation is needed. Our ligand-binding experiments using SPR and DSF did not support previous *in silico* predictions of direct binding between famotidine and SARS-CoV-2 proteases. We further used an array of experimental approaches to show that famotidine had no effect on SARS-CoV-2 protease function or generally on viral replication. It must be noted that since the clinical studies correlated putative clinical benefit with the use of higher doses of famotidine, we tested famotidine at significantly higher *in vitro* concentrations than the peak plasma concentrations (0.5–2μM) achieved in the blood of patients in both clinical studies ^6,7^Our data strongly suggest that the probable clinical benefit of famotidine likely arises independently of an antiviral mechanism of action.

COVID-19 complications are associated with a severe pro-inflammatory response in the lungs of infected patients^28^. The “cytokine storm” as a result of inflammation is a key pathognomonic feature of COVID-19 and the main contributor to respiratory failure and mortality^29^. Severe COVID-19 cases are characterized by pulmonary infiltration and extensive pulmonary edema, causing exudation of inflammatory cells in the alveolar space, resulting in extensive pulmonary consolidation leading to pneumonia and adult respiratory distress syndrome (ARDS)^30–33^. The results of the two famotidine-related COVID-19 clinical reports, when taken together ^6,7^ suggest that famotidine likely helps with mitigating moderate to severe respiratory symptoms ranging from shortness of breath to intubation. Our data does not rule out the possibility that famotidine related improvements in COVID-19 patients are through an antiinflammatory action. For example, the development of the cytokine storm in COVID-19 patients is characterized by elevation of pro-inflammatory type I cytokines, which are secreted from a variety of cells such as polymorphonuclear cells, natural killer cells, and endothelial cells, etc^29^. It is therefore conceivable that famotidine-related benefit in managing respiratory symptoms may be due to an anti-inflammatory mechanism of action.

It is noteworthy that H2R, the established molecular target of famotidine, is involved in the activation of several mediators of the adaptive immune response, such as Th1 lymphocytes, which are implicated in pro-inflammatory cytokine production^34^. Histamine, the H2R ligand, also regulates bronchoconstriction, airway inflammation, and vasodilation^34^. Mast cells are a major source of histamine and their activation has been reported following viral infections of the respiratory tract^35–37^. Therefore, Mast cells may represent an underappreciated source of pro-inflammatory cytokine release in COVID-19 patients^35^. A better understanding of the role of the H2R pathway in COVID-19 will help elucidate the molecular details of how famotidine reduces the disease severity.

Our study redirects the mechanism behind the potential beneficial effect of famotidine, away from an antiviral effect to likely an anti-inflammatory action in COVID-19 patients. Given that there is an ongoing randomized clinical trial (NCT04370262), our results may assist the investigators in reshaping their interventional study to include inflammation-related outcomes. Also, it should be noted that while famotidine is one of the relatively safer drugs, its use is not without risk^38–40^, especially in elderly patients (a high-risk population for COVID-19), in which famotidine use has been associated with CNS complications^41^. Provided the ongoing clinical trial yields promising results, further investigation of famotidine and its safety profile in different age brackets will be needed before the drug can be used, most likely as part of a combination therapy, for COVID-19 disease management.

## Author contributions

Conceptualization: AHM; data curation and analysis: HL, AB, SB, MS, AHM, methodology and investigation: ML, HL, DYC, SDG, SB, AM, SDW, AO, FD MS, AHM; writing (original draft): AHM; writing (review and editing): AHM, MS, FD, SDW, HL, SB

## Supplementary Figures

**Table S1:**
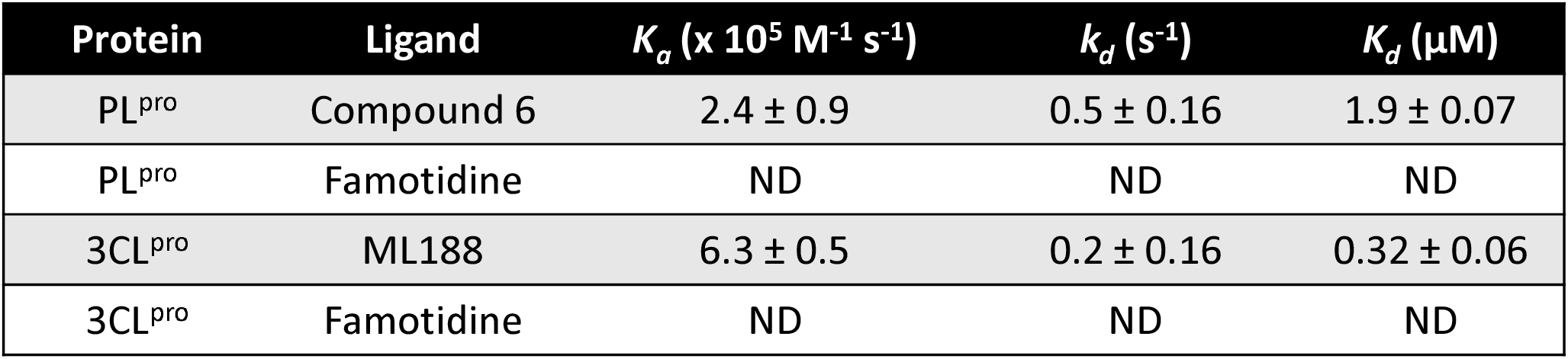
Summary of the kinetic and equilibrium dissociation constants determined by SPR at 10 °C. The *K_d_* for *compound 6* and *ML188* were determined from the binding isotherms shown in **Figure 2. B & D** (in the main text). Famotidine and *compound 6* were tested up to a maximal dose of 50 mM and *ML188* up to 5 mM. No binding was detected for famotidine to either protein. The *K_d_* for the tool compounds was determined from analysis of the binding isotherms (**B & D**). Standard deviation from the mean was determined from triplicate experiments.

**Figure S1:**
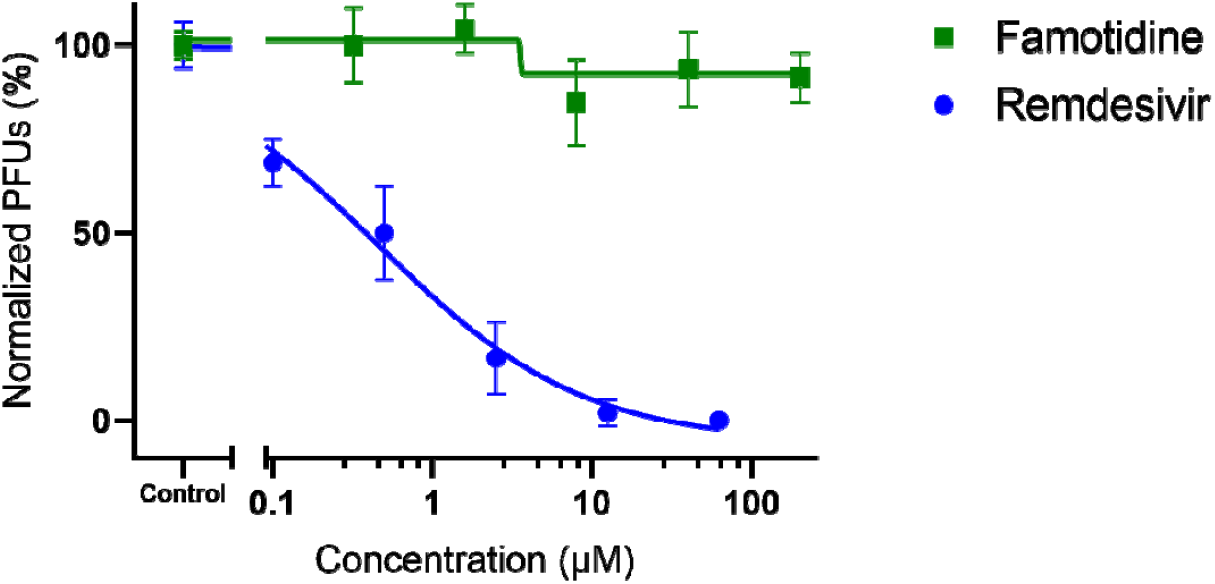
Antiviral effect of famotidine on virus yield (plaque formation) in Vero E6 cells. Remdisivir (blue) produced a dose-dependent antiviral effect when tested at a range of 0.1 μM and 62.5 μM concentrations. Famotidine (green) produced no effect on limiting virus plaque formation when tested between the ranges of 0.32 μM and 200 μM. Each point represents the mean ± standard deviation of triplicates.

**Figure S2.**
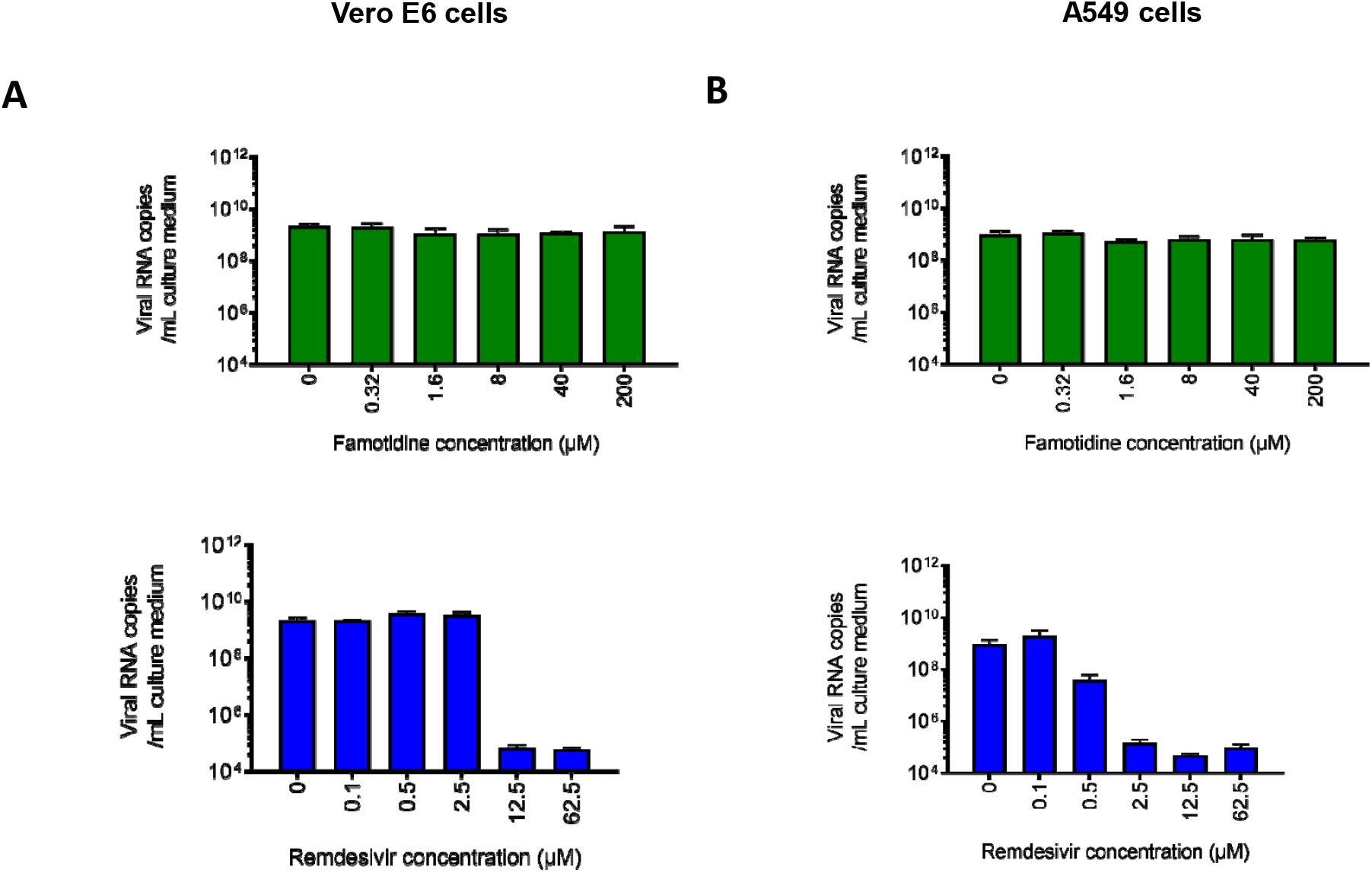
Antiviral activity of famotidine in Vero E6 and human lung A549 cells assessed by qRT-PCR. Quantitative real-time PCR was used to determine SARS-CoV-2 RNA with primers for the viral envelope (E) gene. Viral mRNA copies/mL of culture medium are shown when infected cells were grown in the presence of a range of famotidine (green) and remdesivir (blue) concentrations. While remdesivir exerts a dose-dependent effect on suppressing virus replication as seen by reduction in viral RNA copies, famotidine showed no effect in either Vero E6 or human lung A549 cells.

**Figure S3.**
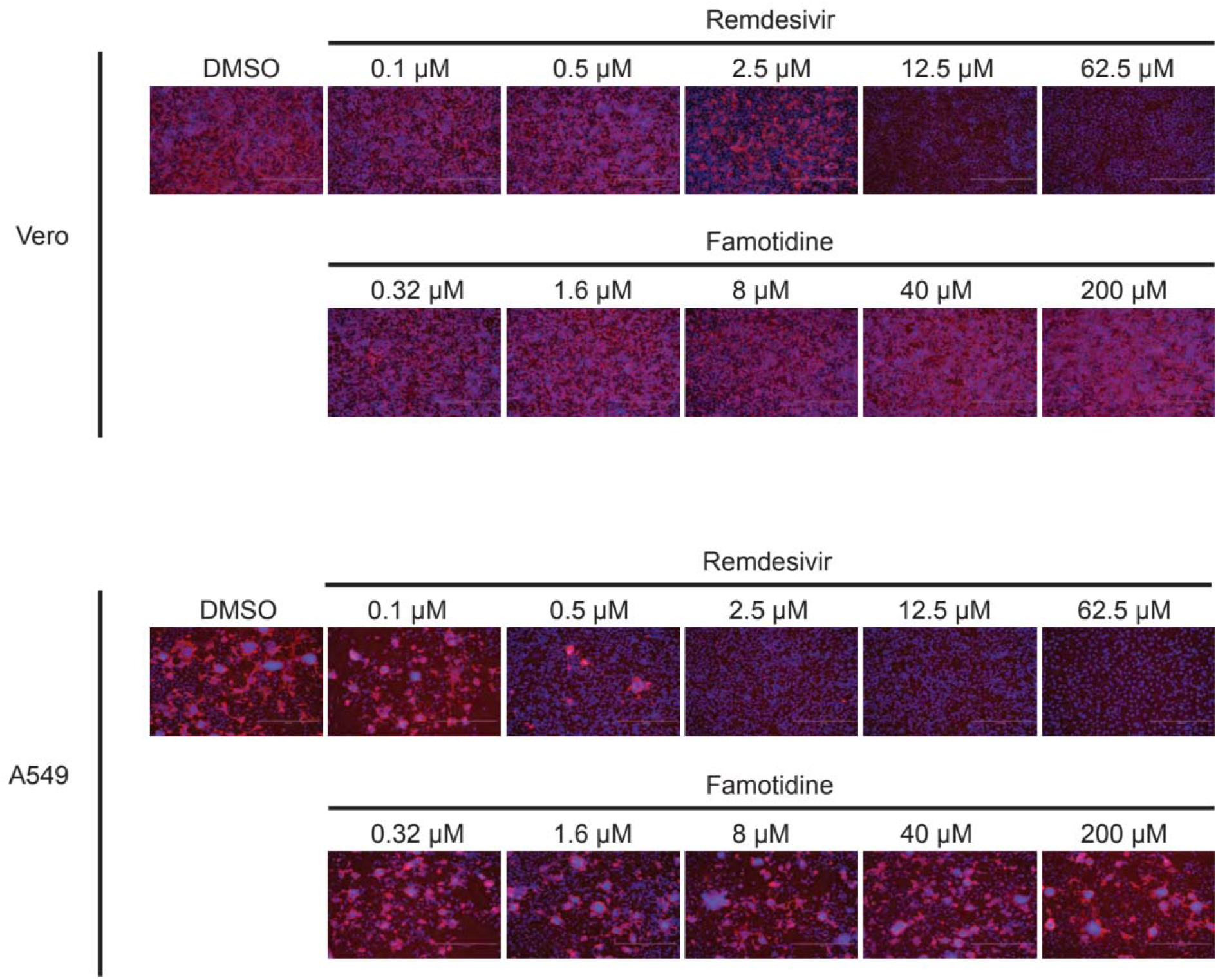
Immunofluorescence staining of SARS-CoV-2 infected Vero E6 and human lung A549 cells in the presence of famotidine. Virus infected cells were fixed with paraformaldehyde and stained with an antibody directed to the virus N protein.

## REFERENCES

1. Harrison C. Coronavirus puts drug repurposing on the fast track. Nat Biotechnol 2020;38:379–81.

2. Beigel JH, Tomashek KM, Dodd LE, et al. Remdesivir for the Treatment of Covid-19 - Preliminary Report. N Engl J Med 2020.

3. Boulware DR, Pullen MF, Bangdiwala AS, et al. A Randomized Trial of Hydroxychloroquine as Postexposure Prophylaxis for Covid-19. N Engl J Med 2020.

4. Cao B, Wang Y, Wen D, et al. A Trial of Lopinavir-Ritonavir in Adults Hospitalized with Severe Covid-19. N Engl J Med 2020;382:1787–99.

5. Keithley JK. Histamine H2-receptor antagonists. Nurs Clin North Am 1991;26:361–73.

6. Freedberg DE, Conigliaro J, Wang TC, et al. Famotidine Use is Associated with Improved Clinical Outcomes in Hospitalized COVID-19 Patients: A Propensity Score Matched Retrospective Cohort Study. Gastroenterology 2020.

7. Janowitz T, Gablenz E, Pattinson D, et al. Famotidine use and quantitative symptom tracking for COVID-19 in non-hospitalised patients: a case series. Gut 2020.

8. Wu C, Liu Y, Yang Y, et al. Analysis of therapeutic targets for SARS-CoV-2 and discovery of potential drugs by computational methods. Acta Pharm Sin B 2020.

9. Shaffer L. 15 drugs being tested to treat COVID-19 and how they would work. Nat Med 2020.

10. Zhang L, Lin D, Sun X, et al. Crystal structure of SARS-CoV-2 main protease provides a basis for design of improved alpha-ketoamide inhibitors. Science 2020;368:409–12.

11. Muramatsu T, Takemoto C, Kim YT, et al. SARS-CoV 3CL protease cleaves its C-terminal autoprocessing site by novel subsite cooperativity. Proc Natl Acad Sci U S A 2016;113:12997–3002.

12. Tomar S, Johnston ML, St John SE, et al. Ligand-induced Dimerization of Middle East Respiratory Syndrome (MERS) Coronavirus nsp5 Protease (3CLpro): IMPLICATIONS FOR nsp5 REGULATION AND THE DEVELOPMENT OF ANTIVIRALS. J Biol Chem 2015;290:19403–22.

13. Baez-Santos YM, Mielech AM, Deng X, Baker S, Mesecar AD. Catalytic function and substrate specificity of the papain-like protease domain of nsp3 from the Middle East respiratory syndrome coronavirus. J Virol 2014;88:12511–27.

14. Jacobs J, Grum-Tokars V, Zhou Y, et al. Discovery, synthesis, and structure-based optimization of a series of N-(tert-butyl)-2-(N-arylamido)-2-(pyridin-3-yl) acetamides (ML188) as potent noncovalent small molecule inhibitors of the severe acute respiratory syndrome coronavirus (SARS-CoV) 3CL protease. J Med Chem 2013;56:534–46.

15. Jacobs J, Zhou S, Dawson E, et al. Discovery of non-covalent inhibitors of the SARS main proteinase 3CLpro. Probe Reports from the NIH Molecular Libraries Program. Bethesda (MD) 2010.

16. Ratia K, Pegan S, Takayama J, et al. A noncovalent class of papain-like protease/deubiquitinase inhibitors blocks SARS virus replication. Proc Natl Acad Sci U S A 2008;105:16119–24.

17. Lu IL, Mahindroo N, Liang PH, et al. Structure-based drug design and structural biology study of novel nonpeptide inhibitors of severe acute respiratory syndrome coronavirus main protease. J Med Chem 2006;49:5154–61.

18. Lindner HA, Fotouhi-Ardakani N, Lytvyn V, Lachance P, Sulea T, Menard R. The papain-like protease from the severe acute respiratory syndrome coronavirus is a deubiquitinating enzyme. J Virol 2005;79:15199–208.

19. Barretto N, Jukneliene D, Ratia K, Chen Z, Mesecar AD, Baker SC. The papain-like protease of severe acute respiratory syndrome coronavirus has deubiquitinating activity. J Virol 2005;79:15189–98.

20. Freitas BT, Durie IA, Murray J, et al. Characterization and Noncovalent Inhibition of the Deubiquitinase and deISGylase Activity of SARS-CoV-2 Papain-Like Protease. ACS Infect Dis 2020.

21. Sun C, Li Y, Yates EA, Fernig DG. SimpleDSFviewer: A tool to analyze and view differential scanning fluorimetry data for characterizing protein thermal stability and interactions. Protein Sci 2020;29:19–27.

22. Ogando NS, Dalebout TJ, Zevenhoven-Dobbe JC, et al. SARS-coronavirus-2 replication in Vero E6 cells: replication kinetics, rapid adaptation and cytopathology. J Gen Virol 2020.

23. Corman VM, Landt O, Kaiser M, et al. Detection of 2019 novel coronavirus (2019- nCoV) by real-time RT-PCR. Euro Surveill 2020;25.

24. van Hemert MJ, van den Worm SH, Knoops K, Mommaas AM, Gorbalenya AE, Snijder EJ. SARS-coronavirus replication/transcription complexes are membrane-protected and need a host factor for activity in vitro. PLoS Pathog 2008;4:e1000054.

25. Kung JE, Jura N. Structural Basis for the Non-catalytic Functions of Protein Kinases. Structure 2016;24:7–24.

26. Xie X, Muruato AE, Zhang X, et al. A nanoluciferase SARS-CoV-2 for rapid neutralization testing and screening of anti-infective drugs for COVID-19. bioRxiv 2020.

27. Palacio-Rodriguez K, Lans I, Cavasotto CN, Cossio P. Exponential consensus ranking improves the outcome in docking and receptor ensemble docking. Sci Rep 2019;9:5142.

28. Tay MZ, Poh CM, Renia L, MacAry PA, Ng LFP. The trinity of COVID-19: immunity, inflammation and intervention. Nat Rev Immunol 2020;20:363–74.

29. Ye Q, Wang B, Mao J. The pathogenesis and treatment of the ‘Cytokine Storm’ in COVID-19. J Infect 2020;80:607–13.

30. Carsana L, Sonzogni A, Nasr A, et al. Pulmonary post-mortem findings in a series of COVID-19 cases from northern Italy: a two-centre descriptive study. Lancet Infect Dis 2020.

31. Bhatraju PK, Ghassemieh BJ, Nichols M, et al. Covid-19 in Critically Ill Patients in the Seattle Region - Case Series. N Engl J Med 2020;382:2012–22.

32. Tian J, Yuan X, Xiao J, et al. Clinical characteristics and risk factors associated with COVID-19 disease severity in patients with cancer in Wuhan, China: a multicentre, retrospective, cohort study. Lancet Oncol 2020.

33. Yang K, Sheng Y, Huang C, et al. Clinical characteristics, outcomes, and risk factors for mortality in patients with cancer and COVID-19 in Hubei, China: a multicentre, retrospective, cohort study. Lancet Oncol 2020.

34. Thangam EB, Jemima EA, Singh H, et al. The Role of Histamine and Histamine Receptors in Mast Cell-Mediated Allergy and Inflammation: The Hunt for New Therapeutic Targets. Front Immunol 2018;9:1873.

35. Marshall JS, Portales-Cervantes L, Leong E. Mast Cell Responses to Viruses and Pathogen Products. Int J Mol Sci 2019;20.

36. Zarnegar B, Mendez-Enriquez E, Westin A, et al. Influenza Infection in Mice Induces Accumulation of Lung Mast Cells through the Recruitment and Maturation of Mast Cell Progenitors. Front Immunol 2017;8:310.

37. Hu Y, Jin Y, Han D, et al. Mast cell-induced lung injury in mice infected with H5N1 influenza virus. J Virol 2012;86:3347–56.

38. Kirch W, Halabi A, Linde M, Santos SR, Ohnhaus EE. Negative effects of famotidine on cardiac performance assessed by noninvasive hemodynamic measurements. Gastroenterology 1989;96:1388–92.

39. Lee YC, Wang CC. Famotidine-induced retinopathy. Eye (Lond) 2006;20:260–3.

40. Kallal SM, Lee M. Thrombotic thrombo-cytopenic purpura associated with histamine H2-receptor antagonist therapy. West J Med 1996;164:446–8.

41. Cantu TG, Korek JS. Central nervous system reactions to histamine-2 receptor blockers. Ann Intern Med 1991;114:1027–34.

